# A systematic evaluation revealed that detecting translated non-canonical ORFs from ribosome profiling data remains challenging

**DOI:** 10.1101/2022.12.11.520003

**Authors:** Tianyu Lei, Yue Chang, Chao Yao, Hong Zhang

## Abstract

Non-canonical open reading frames (ORFs), which are ORFs that are not included in reference genome annotations, are gaining more and more research interest in recent years. While vast numbers of non-canonical ORFs have been identified with ribosome profiling (Ribo-Seq) by various state-of-the-art computational methods, the performance of these methods has not been assessed systematically. To this end, we evaluated the four most popular methods for translated non-canonical ORF prediction using various public datasets. We found that non-canonical ORFs predicted by different methods differ substantially and are not saturated at typical sequence depths. Furthermore, the precision and accuracy of all four methods are not satisfactory, especially for ORFs with near-cognate start codons. Based on these results, we suggest that improved sequence depth, biological repetitions, and translation initiation site profiling should be considered to obtain a high-quality catalog of translated non-canonical ORFs in future studies.

## Introduction

A fundamental step in dissecting the genetic information stored in an organism’s genome is to identify all the coding elements transcribed to messenger RNAs (mRNAs) and further translated to various proteins. In eukaryotes, mRNAs are believed to contain a single ORF as the coding sequence (CDS) due to the cap-dependent mechanism of translation initiation [1, 2]. However, it was observed decades ago that ORFs initiated from alternative start codons in 5’ untranslated regions (UTRs) could fine-tune mRNA translation and stability in *cis* [3–5]. Furthermore, some RNAs presumed to be non-coding are found to encode novel functional peptides by recent studies[6–10]. While such non-canonical ORFs play essential roles in diverse biological processes[11–15], they were often overlooked in traditional studies due to difficulties in detection.

This situation has gradually changed since the advent of ribosome profiling (Ribo-Seq)[16]. During Ribo-Seq, ribosome-protected fragments are enriched and sequenced, which enables genome-wide profiling of translational activities at nucleotide resolution [16]. Since then, Ribo-Seq has been adapted to characterize translation mechanisms with decreasing sample requirements[17–19] or from different facets such as 40S subunits [20], initiating ribosomes[21, 22], active ribosomes [23], and stalled ribosomes [24, 25]. Based on the characteristic distribution of ribosome-protected fragments (RPFs) in Ribo-Seq, such as enrichment in translated regions and three-nucleotide periodicity [26], many computational methods have been developed to detect translated ORFs from Ribo-Seq data [27, 28]. The continuing development of Ribo-Seq techniques and related computational methods leads to the identification of tens of thousands of non-canonical ORFs in UTRs of mRNAs and long non-coding RNAs (lncRNAs) in different species [29, 30]. Specialized databases, including SmProt[31], OpenProt[32], RPFdb[33], and sORFs.org[34], have been created to catalog the vast amount of non-canonical ORFs identified by different methods.

However, methods used for predicting non-canonical ORFs differ study by study, and there is no consensus on best practices. For example, early methods rely on users to supply the list of potentially translated ORFs for assessment [35–37]. Differences in criteria used for candidate ORF preparation could introduce biases in the final results. While many non-canonical ORFs found by independent studies use near-cognate start codons for initiation [38], some methods consider none or only subsets of near-cognate codons when predicting translated ORFs [35, 39–41]. Besides, some methods require Ribo-Seq conducted with different translation inhibitors [41] or RNA-Seq of the same samples [39]. Differences in workflows of translated ORF prediction might lead to discrepant results and impede the further advancement of this field and related communications. As non-canonical ORFs are gaining more and more interest, we anticipate that a systematic evaluation of computational methods for translated non-canonical ORFs discovery from Ribo-Seq data would be helpful for future studies on the diversity, functions, and evolution of non-canonical ORFs.

To close this gap, here we take the first step by evaluating non-canonical ORFs identified from conventional methods by several popular methods, including PRICE [42], Ribo-TISH [43], RiboCode [44], and RibORF [45]. We examined whether non-canonical ORFs identified in a sample are saturated for different methods and evaluated their performance in terms of accuracy and precision. Based on these results, we provided suggestions on best practices of non-canonical ORF discovery.

## Results

### Choice of software and benchmark datasets

To maximize the generalizability of this study, we focused on methods that only require conventional Ribo-Seq data as input since matched translation initiation sequencing (TI-Seq) data are unavailable for most public datasets. Furthermore, we excluded methods that do not allow near-cognate start codons that have been shown to initiate translation by various evidence [46, 47]. Among methods meeting these criteria, we chose to benchmark PRICE, RiboCode, RibORF, and Ribo-TISH, since they have seen wide adoption in recent large-scale studies [13, 14, 48] or in building databases of translated non-canonical ORFs[49–51].

To evaluate the reliability of predicted non-canonical ORFs, we obtained raw data from 21 Ribo-Seq libraries that have matched TI-Seq from previous studies [12, 22, 41] (Table S1). This dataset encompassed various cell lines or tissues from humans or mice, which could alleviate potential selection biases. In addition, the dataset included two samples with three biological replicates, making it possible to evaluate the precision of ORF predictions. We predicted translated ORFs using all four methods for each library and processed the prediction results with a unified pipeline (Methods). The total number of translated non-canonical ORFs in the same sample varies widely across different prediction methods (Fig. 1A). Overall, RibORF predicted the highest numbers of non-canonical ORFs, while PRICE predicted the least numbers in most samples.

**Figure 1.**
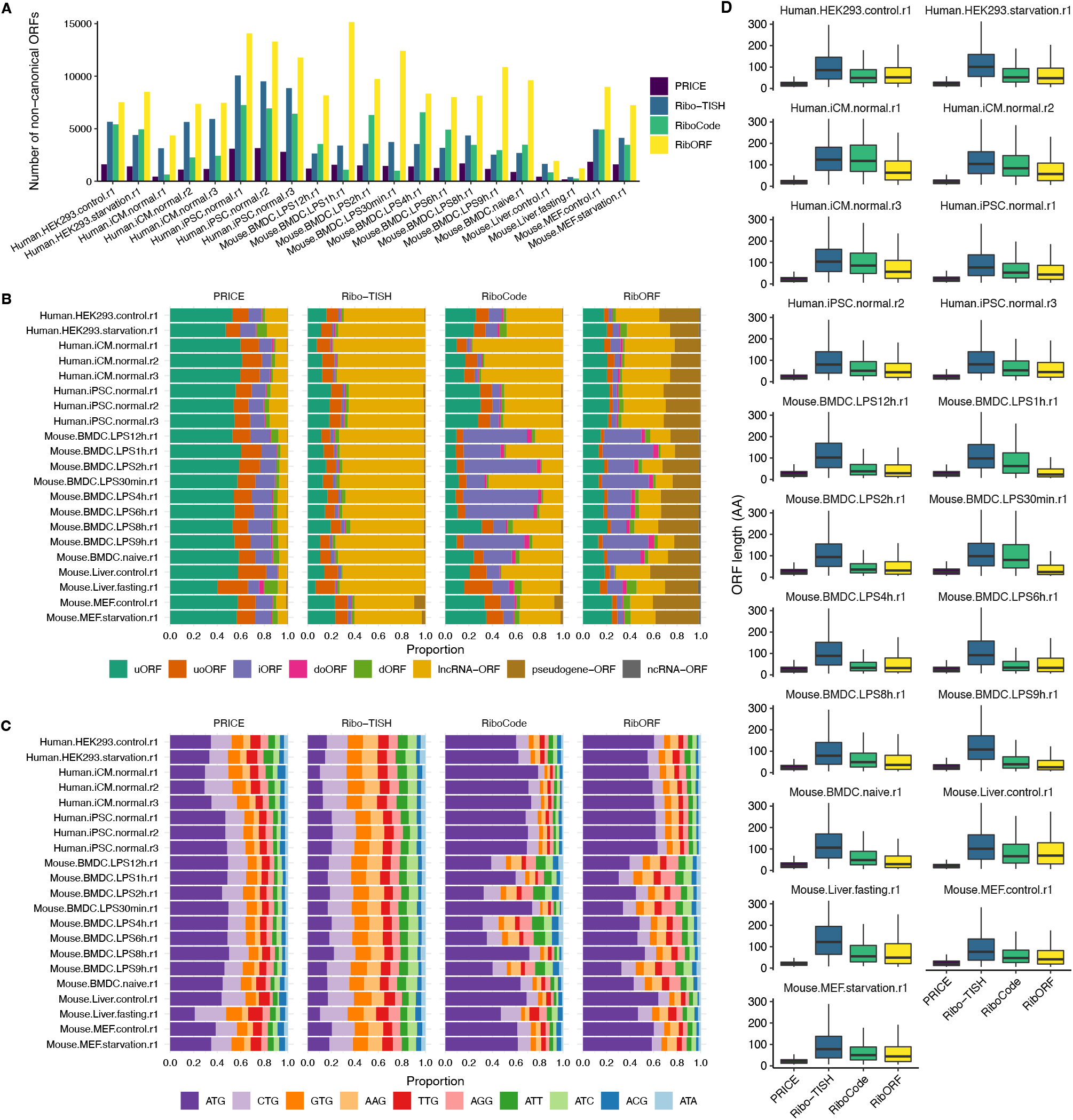
Comparison of non-canonical ORFs detected by different methods. (**A**) The total number of translated non-canonical ORFs detected by different methods in each sample. (**B**, **C**) Fraction of ORFs belonging to different types (**B**) or with different start codons (**C**) for non-canonical ORFs presented in **A**. (**D**) ORF length distributions for non-canonical ORFs are presented in **A**. Stop codons were included when calculating ORF lengths in amino acids (AA).

### Composition of predicted translated non-canonical ORFs

Then following a recent community proposal for translated ORF annotation [38], we classified the non-canonical ORFs into different categories according to the type of transcripts where they reside and their positions relative to main ORFs. Specifically, non-canonical ORFs in protein-coding transcripts are categorized into upstream ORFs (uORFs), upstream-overlapping ORFs (uoORFs), internal ORFs (iORFs), downstream-overlapping ORFs (doORFs) and downstream ORFs (dORFs), while those located in lncRNAs, pseudogene transcripts, and the remaining non-coding RNAs (ncRNAs) are categorized into lncRNA-ORFs, pseudogene-ORFs, and ncRNA-ORFs, respectively (Fig. S1). Here, we included two new categories, pseudogene-ORFs and ncRNA-ORFs, as they are discovered by all four methods and might have novel functions deserving further study.

We found that the composition of non-canonical ORFs predicted by each method differs substantially from those predicted by the other three methods (Fig.1B). Non-canonical ORFs identified by PRICE are mostly (89.4±3.8%, mean±SD) located in mRNAs, while those identified by Ribo-TISH are dominated by lncRNA-ORFs. Besides, RibORF predicted many (29.3±6.0%) pseudogene-ORFs, which are rare among ORFs predicted by the other three methods. Consistent with the cap-dependent mechanism of translation initiation[1, 2], most predicted non-canonical ORFs within mRNAs are uORFs or uoORFs (Fig. 1B). Noteworthy, both RiboCode and RibORF predicted substantially more internal ORFs (41.1±20.1% and 28.5±10.5%, respectively) in mouse bone marrow-derived dendritic cells (BMDC) than the other samples, which might be caused by specialized translational regulation in BMDC.

Next, we examined the start codon usage among non-canonical ORFs predicted by different methods. Except for Ribo-TISH, the most frequently used start codon among non-canonical ORFs predicted by the other three methods is AUG, followed by CTG, GTG, AAG, and TTG, while the other near-cognate start codons are used to a less extent. For non-canonical ORFs predicted by Ribo-TISH, the most frequent start codon is CTG rather than ATG (Fig. 1C). Across different samples, the start codon usage among non-canonical ORFs predicted by PRICE and Ribo-TISH are relatively stable, while it is more variable in the prediction results of RiboCode and RibORF. Since non-canonical ORFs are typically shorter, we also compared length distributions of non-canonical ORFs predicted by different methods. We found that non-canonical ORFs predicted by Ribo-TISH are much longer, while those predicted by PRICE are overall the shortest (Fig. 1D). This trend is evident even among ORFs of the same type (Fig. S2). To summarize, non-canonical ORFs predicted by different methods showed substantial variation in total number, composition, start codon usage, and length distribution, which further underscores the necessity of a systematic evaluation of their prediction results.

### Whether numbers of predicted non-canonical ORFs are saturated?

As the total number of predicted translated non-canonical ORFs varied widely across different methods, a natural question is whether translated non-canonical ORFs identified by different methods are saturated at the current sequencing depth. To address this issue, we mocked up two ultra-deep datasets by combining the three human iPSC libraries and the naïve or LPS-treated mouse BMDC libraries, respectively. Then we randomly sampled different fractions (10% to 90% with a step size of 10%) of raw reads from the two datasets and predicted translated non-canonical ORFs using all four methods with the down-sampled or entire datasets. Finally, we plotted the total number of translated non-canonical ORFs against fractions of reads used for prediction. We envisaged a plateau if the total number of predicted non-canonical ORFs was saturated at a certain point. However, in both datasets, we failed to observe any plateau phase (Fig. 2A). We also stratified non-canonical ORFs by type or start codon identity and observed similar trends among ORFs of different groups (Fig. S3). This suggests that none of the two datasets approximated the saturation sequencing depth for any method.

**Figure 2.**
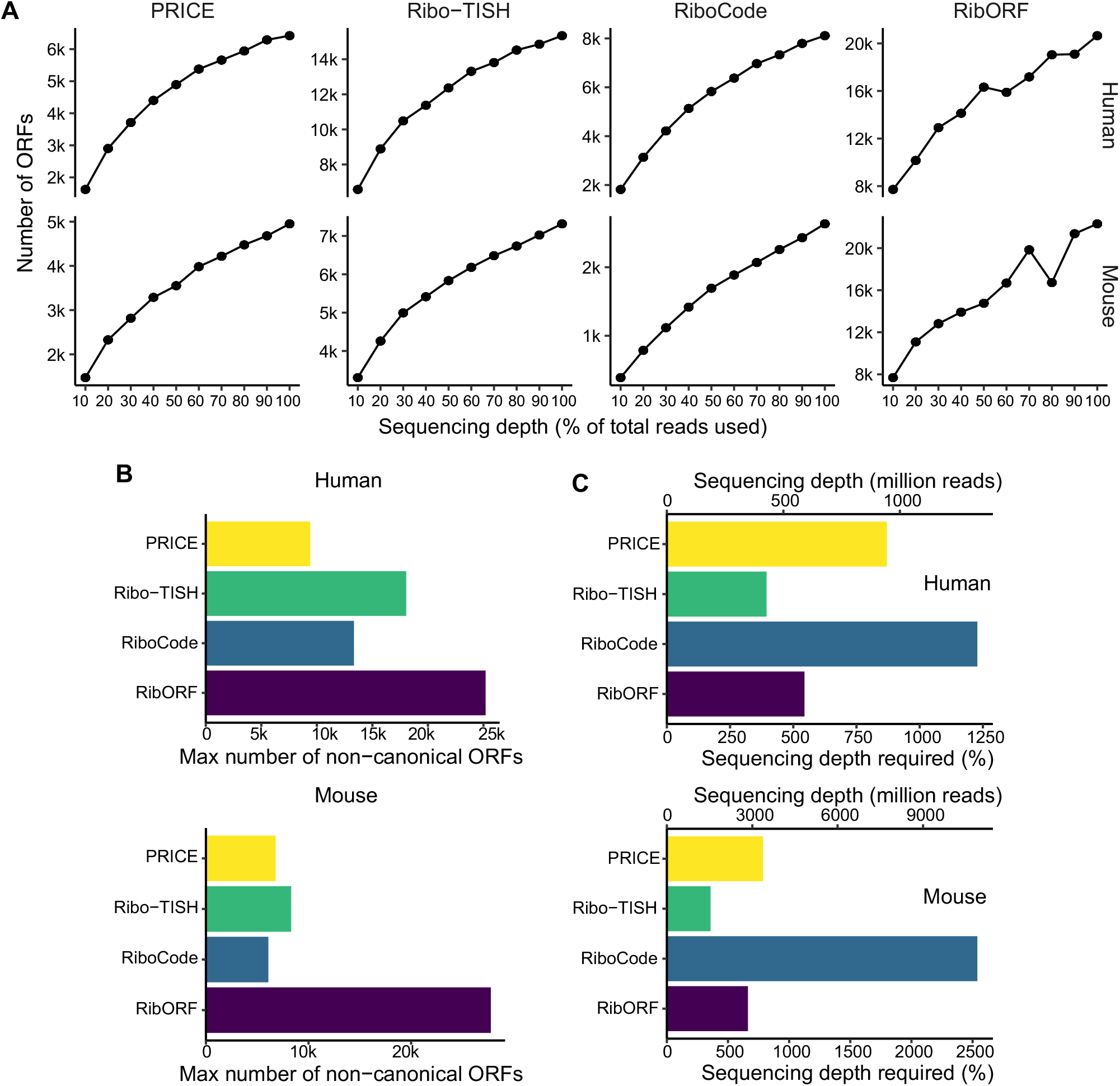
Saturation analysis of the total number of predicted non-canonical ORFs. (**A**) The total number of detected non-canonical ORFs when different fractions of RPF reads were sampled randomly for the analysis. (**B**) The maximum number of non-canonical ORFs can be identified from human or mouse datasets. (**C**) Sequencing depth (in million reads or percent of current depth) required to discover 95% of the maximum number of non-canonical ORFs.

As the total number of predicted non-canonical ORFs increase with reduced rates (Fig. 2A), it is expected to saturate at increased sequencing depth. Therefore, we fitted a non-linear model of the number of predicted ORFs against the sequencing depth for each method, and estimated the maximum number of non-canonical ORFs that could be identified asymptotically (Fig. 2B). Then we calculated the saturation sequencing depth as the depth when 95% of the asymptotic maximum number of non-canonical ORFs could be identified. For human (mouse) data, we estimated the saturation sequencing depth as 3.9~12.3(3.6~25.4) times of the current sequencing depth depending on methods, which translates to 427.7~1332.2 (1534.9~10904.9) million reads (Fig. 2C). With increasing sequencing depth, the results of Ribo-TISH were estimated to saturate first, followed by PRICE and RibORF, and RiboCode requires the highest sequencing depth (Fig. 2C). In summary, the current sequencing depth is too shallow to recovered all potentially translated non-canonical ORFs in a sample for existing methods.

### Comparison of accuracy

Due to the lacking of appropriate ground truth datasets, we cannot benchmark the performance of different methods in standard ways. However, a more accurate method should predict a higher proportion of ORFs supported by independent datasets. Therefore, we evaluated the accuracy of different methods by examining the proportion of translated non-canonical ORFs predicted by each method that can be recovered from TI-Seq data of matched samples. We first predicted non-canonical ORFs independently from TI-Seq libraries only using Ribo-TISH. Samples with < 500 non-canonical ORFs identified from TI-Seq data were excluded due to limited statistical power. Then we measured the accuracy as the proportion of non-canonical ORFs identified from conventional Ribo-Seq data by a method that Ribo-TISH also discovered in the matched TI-Seq data. We found that the prediction accuracy decreased in the order of PRICE > RiboCode > Ribo-TISH > RibORF (Fig. 3A). Noteworthy, we found that the overall accuracy of all four software was not very high, and none of them exceeded 30% in any sample.

**Figure 3.**
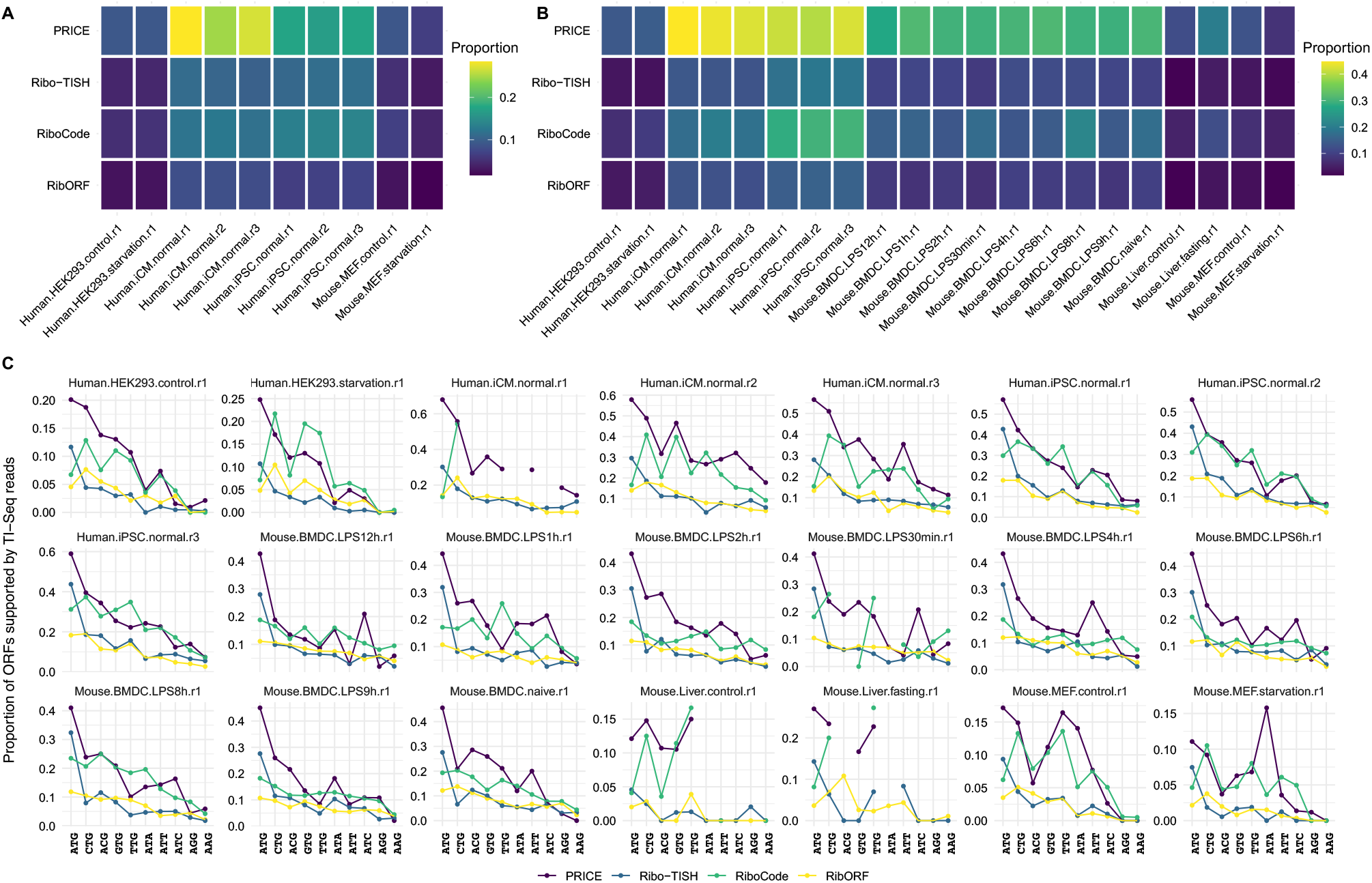
Accuracy of translated non-canonical ORFs predicted by different methods. (**A**) Proportions of non-canonical ORFs identified from conventional Ribo-Seq data by different methods that were also identified with TI-Seq from matched samples by Ribo-TISH. (**B**) Proportions of non-canonical ORFs with RPF peaks around start codons in TI-Seq of matched samples. (**C**) Similar to **B** but stratified by the identity of start codons.

To exclude potential biases introduced during ORF prediction from TI-Seq data, we further calculated the genome-wide P-site coverage for each TI-Seq library. Then we examined whether there are TI-Seq read peaks around the start codons of non-canonical ORFs predicted from conventional Ribo-Seq data (Methods). Across the four methds, we found that PRICE and RiboCode predicted higher proportions of non-canonical ORFs supported by TI-Seq reads (Fig. 3B). Among non-canonical ORFs with different start codons, a higher fraction of ATG ORFs have start codons supported by TI-Seq reads (Fig. 3C). Among non-canonical ORFs of different types, we found that uORFs and uoORFs have higher proportions of start codons supported by TI-Seq reads than the other categories of non-canonical ORFs, especially pseudogene-ORFs (Fig. S4A). Contrary to our expectation, start codons of shorter ORFs are more likely to be supported by TI-Seq reads (Fig. S4B). A possible explanation is that the criteria used to identify translated ORFs are tailored for short ones, which might be too loose for longer ones. In summary, non-canonical ORFs predicted by PRICE and RiboCode are more accurate than those predicted by Ribo-TISH or RibORF. However, the accuracy of all four methods remains relatively low, especially for ORFs with near-cognate start codons.

### Comparison of precision

A precise method should produce consistent prediction results across different biological replicates of the same sample. In both human induced cardiomyocytes (iCM) and induced pluripotent stem cells (iPSC), we found that non-canonical ORFs predicted by RibORF and RiboCode have the highest extent of overlapping among different biological replicates (Fig. 4A). To quantitatively measure the precision of each method, we summarized the similarity of prediction results across different replicates with Sørensen-Dice coefficient (SDC). SDC is similar to the Jaccard index but could be extended to comparisons involving more than two sets, and a higher value of SDC indicates higher similarity. Consistent with the visual expectation, SDC of RiboCode and RibORF is higher than that of PRICE and Ribo-TISH (Fig. 4B). Analysis of additional human and mouse datasets with at least three replicates revealed similar trends (Fig. 4C). After stratifying non-canonical ORFs by type, we found that iORFs, doORFs, dORFs, and pseudogene-ORFs have lower reproducibility than the remaining types of non-canonical ORFs (Fig. S5A). Among non-canonical ORFs with different start codons, we found that the reproducibility of ATG ORFs is much higher than those with near-cognate start codons (Fig. 4D). Across ORFs of different lengths, longer ORFs tend to be consistently identified across different replicates in all methods except PRICE, where short ones have higher similarity across replicates (Fig. S5B).

**Figure 4.**
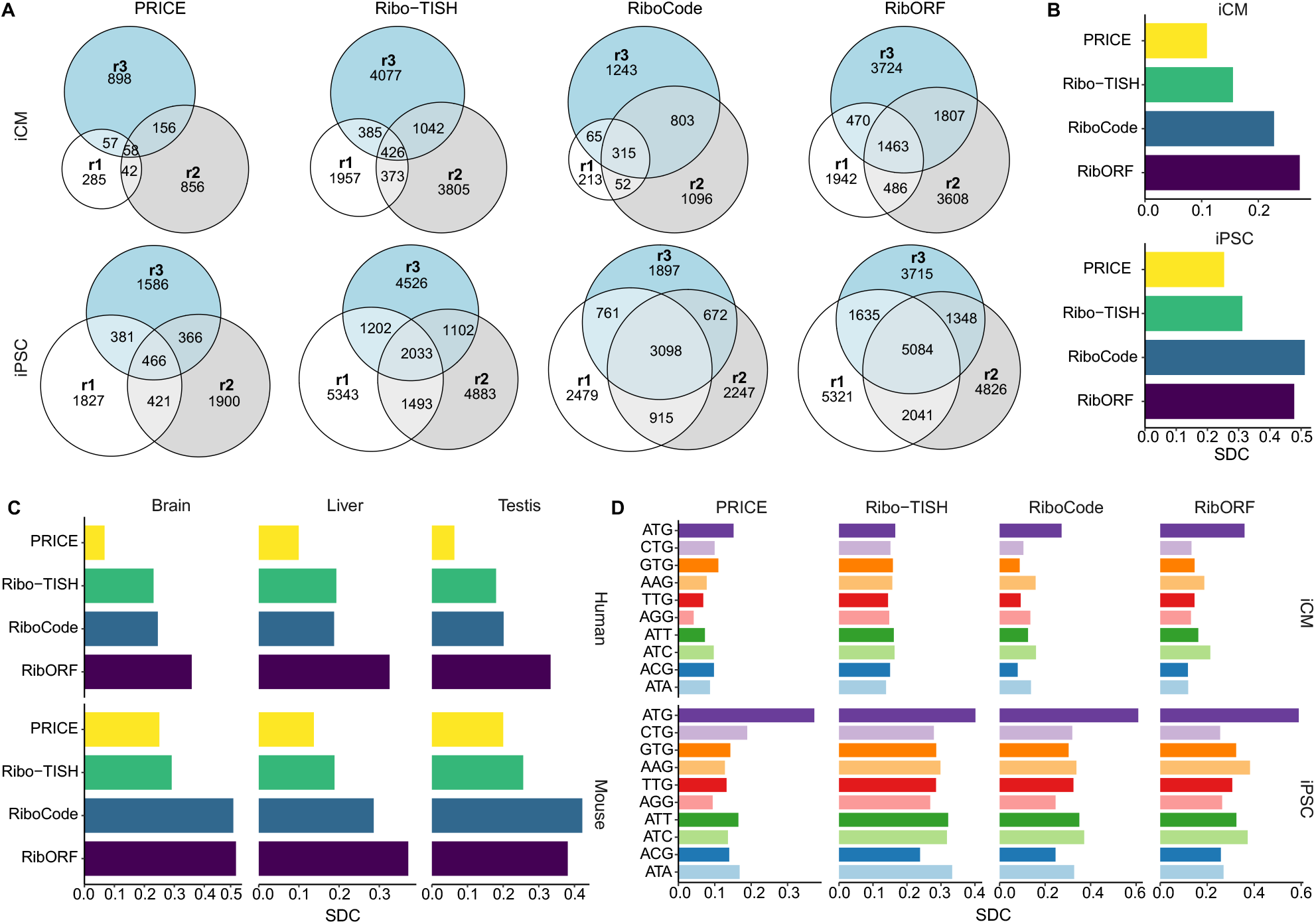
Consistency of predicted non-canonical ORFs across different biological replicates. (**A**) Venn diagram showing overlaps across non-canonical ORFs predicted by the same method across different biological replicates (r1, r2, r3) of the same sample. The number of ORFs in each section was presented. iCM, induced cardiomyocyte; iPSC, induced pluripotent stem cell. (**B**) Similarity of non-canonical ORFs identified from different replicates of human iCM and iPSC libraries as measured by Sørensen-Dice coefficient (SDC). (**C**) Similarity of predicted non-canonical ORFs across different Ribo-Seq replicates of brain, liver, and testis of humans and mice. (**D**) Similar to **B** but stratified by start codon identity.

To further evaluate the consistency of non-canonical ORFs predicted by each method, we examined whether translated non-canonical ORFs identified from down-sampled data can be recovered at higher sequencing depth. We found that a higher proportion of ORFs present in low-depth samples are recovered in high-depth samples by RiboCode and RibORF (Fig. S6), suggesting that their results are more robust than the other two methods. Altogether, our results suggest that RiboCode and RibORF outperform the other two methods in terms of precision when predicting translated non-canonical ORFs. However, the reproducibility of non-ATG ORFs is relatively low in all four methods, which should be properly dealt with in future studies.

### Best practices of translated non-canonical ORF discovery

Given these limitations of existing methods, we explored how to obtain more accurate and precise predictions of translated non-canonical ORFs based on conventional Ribo-Seq data. We first compared the non-canonical ORFs predicted from the same sample by different methods. For non-canonical ORFs identified by each method, many of them are not detected by any of the other three methods (Fig. 5A). After further stratifying ORFs by type or start codon identity. We found that uORFs, uoORFs, and lncRNA-ORFs predicted by different methods are more similar than the other categories (Fig. 5B). In contrast, pseudogene-ORFs and ncRNA-ORFs show the least similarity among different methods and whether they are bona fide translated ORFs requires further verification. Besides, the similarity of ORFs with near-cognate start codons is much lower than that of AUG ORFs predicted by different methods (Fig. 5C). In summary, we found non-canonical ORFs predicted by different methods are highly dissimilar.

**Figure 5.**
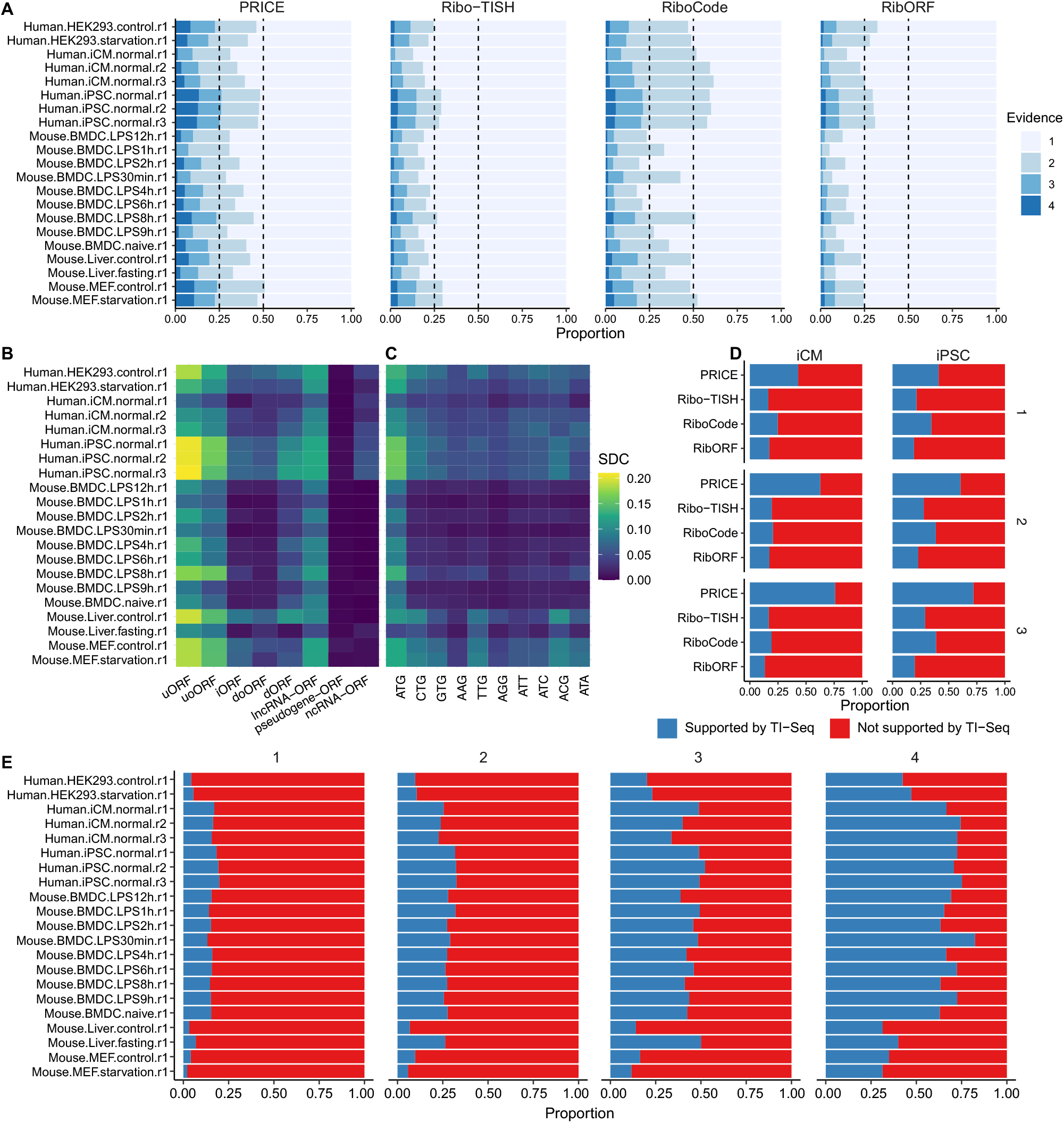
Influences of combining non-canonical ORFs predicted by different methods or from different biological replicates. (**A**) Proportion of ORFs with different evidence levels for non-canonical ORFs predicted by each method. For each ORF, the evidence level measures how many methods identified this ORF in the same library. (**B-C**) The similarity of predicted non-canonical ORFs across different biological replicates after stratifying by ORF types (**B**) or start codon identity (**C**). (**D**) Proportion of non-canonical ORFs supported by peaks of TI-Seq reads for ORFs consistently identified in different numbers of replicate Ribo-Seq libraries of human iCM or iPSC samples. (**E**) Proportion of non-canonical ORFs supported by peaks of TI-Seq reads for ORFs consistently identified by different numbers of methods from the same conventional Ribo-Seq library.

Next, we examined whether combining results from different methods or replicates could improve prediction accuracy or precision. First, considering that non-canonical ORFs predicted from different biological replicates of the same sample varied widely, we examined whether non-canonical ORFs that are consistently identified across different replicates are more likely to be supported by TI-Seq reads. However, in both iCM and iPSC datasets, this trend was only observed for PRICE (Fig. 5D). Second, we examined whether non-canonical ORFs identified by multiple methods have higher accuracy and precision. As expected, ORFs identified by more methods have a higher proportion supported by TI-Seq reads across samples (Fig. 5E). However, in both iCM and iPSC datasets, there is no evidence that ORFs detected in the same library by at least two methods are more likely to be rediscovered across different biological replicates (Fig. S7). These methods may be too sensitive to RPF distribution noise arising from sampling errors or perturbations introduced during library preparation. Taken together, our analysis suggests that focusing on common non-canonical ORFs reproducibly identified across different methods or biological replicates could improve the accuracy of ORF prediction but not precision.

## Discussion

In this study, we systematically evaluated several methods widely adopted by the community to predict translated non-canonical ORFs with Ribo-Seq data. Our analyses revealed limitations of existing methods that preclude us from gaining a deeper understanding of the non-canonical translatome. First, typical sequencing depths in Ribo-Seq datasets are too shallow to detect all potentially translated non-canonical ORFs in a sample. Since non-canonical ORFs are much shorter than CDSs and some might be lowly expressed, such ORFs are prone to be missed by existing methods due to inadequate statistical power. Second, the performance of all four methods evaluated here is not satisfactory in terms of both accuracy and precision, especially for ORFs with near-cognate start codons. We found that less than 30% of non-canonical ORFs predicted from Ribo-Seq data are supported by TI-Seq reads across different samples no matter which method is used. Many non-canonical ORFs cannot be consistently detected across different biological replicates of the same sample. Among the four methods, we found that PRICE and RiboCode generally have higher accuracy, while RiboCode and RibORF have better precision. These limitations suggest that it is more challenging than anticipated to catalog all translated non-canonical ORFs in cells and better methods or workflows should be developed to capture the diversity of non-canonical ORFs.

Based on these results, we also provide practical suggestions on how to identify translated non-canonical ORFs reliably based on Ribo-Seq data. First, Ribo-Seq libraries should be sequenced to reasonably high depths to capture translated ORFs as complete as possible. Second, at least two biological replicates should be performed to reduce the amount of potential spurious ORFs. Third, employing multiple methods to predict translated ORFs could avoid biases of every single method. Finally, and most importantly, matched TI-Seq libraries should be prepared whenever possible for accurate identification of non-canonical ORFs, especially those with near-cognate start codons. Besides, some post hoc filtering steps could be conducted to improve the quality of the prediction results. For example, a recent study filtered candidate ORFs by the periodicity, uniformity and drop-off rate RPF coverage [48]. However, precautions should be taken to avoid introducing potential biases. For example, the uniformity and drop-off rate of P-site distributions are confounded by RPFs originating from other overlapping ORFs. As a result, the above study identified much fewer uoORFs and doORFs than expected [48]. Integrating mass spectrometry data is another promising direction for reliable identification translated ORFs. However, existing experimental or analytical methods should be optimized to discover non-canonical ORF peptides that are usually short and possibly short-lived [38].

Our analyses do have some drawbacks. First, Ribo-Seq data used in this study are mainly from humans and mice, and further studies are required to test whether our observations here could be generalized to other species. Second, a few public datasets with both Ribo-Seq and TI-Seq have biological repetitions, which constrained the breadth of our precision analysis. As more public datasets are available, we hope a more comprehensive benchmark be performed in the future. Third, while Ribo-Seq and TI-Seq libraries analyzed here use RNase I to digest RNA regions that are not protected by ribosomes, some Ribo-Seq libraries use MNase for digestion[52, 53]. How to identify translated ORFs reliably from such libraries deserves further investigation.

Non-canonical ORFs have broadened our understanding of translatome and more and more functional ones are being discovered by various approaches. Our systematic evaluation here will be helpful to future studies in designing Ribo-Seq experiments and choosing the optimal pipeline to predict translated non-canonical ORFs. Besides, our analyses revealed the limitations of current methods and provide insights for developing better methods in the future.

## Methods

### Datasets and preprocessing

Raw reads of Ribo-Seq and matched TI-Seq libraries generated in previous studies were downloaded from Sequence Read Archive [54] (Table S1). Genome sequences and gene annotations for both human (GRCh38) and mouse (GRCm39) were obtained from ENSEMBL Genome Browser[55]. 3’ adaptors were trimmed using cutadapt (v3.7)[56] with parameters “-j8 -m 18 --trim-n -a” when present. rRNA and tRNA reads were filtered out using bowtie2 (v2.4.5) [57] with parameters “-p 8 --local”. The remaining RPF reads were mapped to reference genomes using STAR (v2.7.10a) [58] with parameters “--outSAMtype BAM SortedByCoordinate --quantMode TranscriptomeSAM GeneCounts --outFilterMultimapNmax 1 --outFilterMatchNmin 16 -- alignEndsType EndToEnd --outSAMattributes NH HI AS nM NM MD”. With “--quantMode TranscriptomeSAM”, STAR will not only produce genomic alignments of RPF reads (“genomic bam”) but also translate the genomic alignments into transcript coordinates (“transcriptomic bam”).

### Prediction of translated non-canonical ORFs

For each Ribo-Seq library, all four methods were run to predict translated ORFs. For PRICE (v1.0.3b)[42], we first generated genome indexes for human and mouse using the “indexGenome” module with parameters “-nostar -nokallisto -nobowtie”. Then we ran PRICE with default parameters using genomic bam files as input. We performed multiple testing corrections with the Benjamini-Hochberg method and used 0.05 as the threshold for adjusted *P* values. For RiboCode (v1.2.13)[44], transcript annotations were generated with “prepare_transcripts”. For each transcriptomic bam file, we obtained P-site offsets using “meta_plots” and ran RiboCode with parameters “-A CTG,GTG,TTG,AAG,ACG,AGG,ATA,ATC,ATT,ATG -l no -g”. For naïve mouse BMDC, a slightly lower cutoff of RPFs in frame 0 was used (“--frame-percent” was set to 0.55 instead of the default 0.60). The default adjusted *P* value cutoff of 0.05 was used to call significant hits. For Ribo-TISH (v0.2.7)[43], P-site offsets were estimated using the “quality” module with default parameters and translated ORFs were predicted using the “predict” module with parameters “--alt --framebest”. The default threshold of 0.05 was used for the frame q value reported by Ribo-TISH.

For RibORF (v1.0) [45], RPF reads were mapped using hisat2 (v2.2.1)[59] with default parameters. All possible ORFs with ATG or near-cognate start codons were predicted using “ORFannotate.pl”. The distribution of RPFs of different lengths was estimated with “readDist.pl”. We performed minimal post-filtering by keeping only read lengths where more than 50% of reads have 5’ ends in frame 0 and significantly higher than that in frame 1 and frame 2 (Wilcox signed-rank test, *P* < 0.01). The P-site offset was estimated from the position with the highest RPF 5’ end coverage within 9 ~ 15 nt before annotated start codons. After obtaining the P-site offsets, “offsetCorrect.pl” was run to generate an alignment file of RPF P-sites. Finally, “ribORF.pl” were run with default parameters to predict translated ORFs. The p values reported by RibORF are estimated probabilities that ORFs are translated. To reduce the false positive rate, we used the 10% quantile of p values of canonical ORFs in the default output as the threshold.

The prediction results of all four methods were processed with a unified pipeline. Briefly, we first slimmed down the transcript annotations based on ENSEMBL transcript biotype classification by excluding: (1) transcripts subjected to targeted degradation, (2) transcripts with intron retention, and (3) immunoglobulin or T cell receptor genes. For the predicted translated ORFs, we removed redundant ones and excluded those located in the mitochondrial genome. Then we remapped ORFs to all the compatible transcripts and excluded ORFs whose start or stop codons coincide with those of annotated CDSs, thus leaving only non-canonical ORFs. We classified non-canonical ORFs in protein-coding transcripts into uORFs, uoORFs, iORFs, doORFs, and dORFs following the recently proposed annotation scheme [38]. For ORFs located in lncRNAs or pseudogene transcripts, we classified them into lncRNA-ORFs and pseudogene-ORFs. All the remaining ORFs that are located in other types of ncRNAs are categorized into ncRNA-ORFs. The default minimum ORF length is 5 amino acids or 18nt including the stop codon for both RiboCode and Ribo-TISH, while PRICE and RibORF use 12nt and 6nt, respectively. For fairness, we used18nt (including the stop codon) as the minimum ORF length for all four methods.

### Saturation analysis

To determine whether the total number of predicted ORFs reached saturation level, we created two ultra-high depth libraries by merging the three Ribo-Seq libraries of human iPSCs and all the Ribo-Seq libraries of LPS-treated mouse BMDC samples, respectively. Downsampling of raw RPF reads was performed using the “seq” subcommand from seqtk (v1.3; https://github.com/lh3/seqtk), with parameter “-f” ranging from 0.1 to 0.9 with a step size of 0.1. Both the full data and down-sampled data were processed and used to predict translated ORFs as described above. Nonlinear regressions of the total number of predicted ORFs (y) against the sequencing depth (x) were performed with the formula 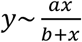 using the “nls” function in R software. The asymptotic maximum number of non-canonical ORFs identifiable is given by parameter *a* in the regression.

### Processing of TI-Seq data

The TI-Seq data were preprocessed and mapped to reference genomes with STAR as described above for Ribo-Seq data. RPFs were assigned to corresponding P-sites with a Random Forest model using the “PSite” package (doi: 10.5281/zenodo.7046270). We retrieved P-site coverage around predicted ORF start codons and considered a start codon as supported by TI-Seq reads if: (1) the first position of the start codon (position 0) has P-site coverage ≥ 5 and (2) total P-site coverage of positions −1, 0, and + 1 is larger than that of positions −4 ~ −2 and positions 2 ~ 4.

To predict translated ORFs with Ribo-TISH[43] from each TI-Seq library, we first ran “quality” module with the parameter “-t”, which indicates that the data is enriched for RPFs around start codons. Then, we ran the “predict” module with parameters “--alt -t” or “--alt --harr -t” for TI-Seq from LTM or harringtonine-treated samples, respectively. Finally, the prediction results were processed with the same pipeline presented above.

### Precision analysis

To quantitatively compare the extent of overlapping across non-canonical ORFs identified from *n* biological replicates, we calculated Sørensen-Dice coefficient (SDC) following a previous study[60]. Let set *S_i_* represent non-canonical ORFs identified from biological replicate *i*. and |*S_i_*| the size of *S_i_*. We calculate SDC as:

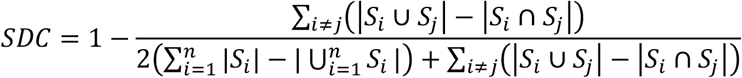

## Data Availability

No new data were generated in this study. Accession code for public datasets used in the analyses is available from Table S1. Custom scripts used for data analysis are available from the corresponding author upon reasonable request.

## Acknowledgments

We thank all the original authors who generated the Ribo-Seq and TI-Seq libraries analyzed in this study and made them publicly available. This work was funded by the National Natural Science Foundation of China (32200433) and the Fundamental Research Funds for the Central Universities (lzujbky-2022-2). We thank the Supercomputing Center of Lanzhou University for providing computational resources.

## Supplementary Tables

**Supplementary Table 1.**
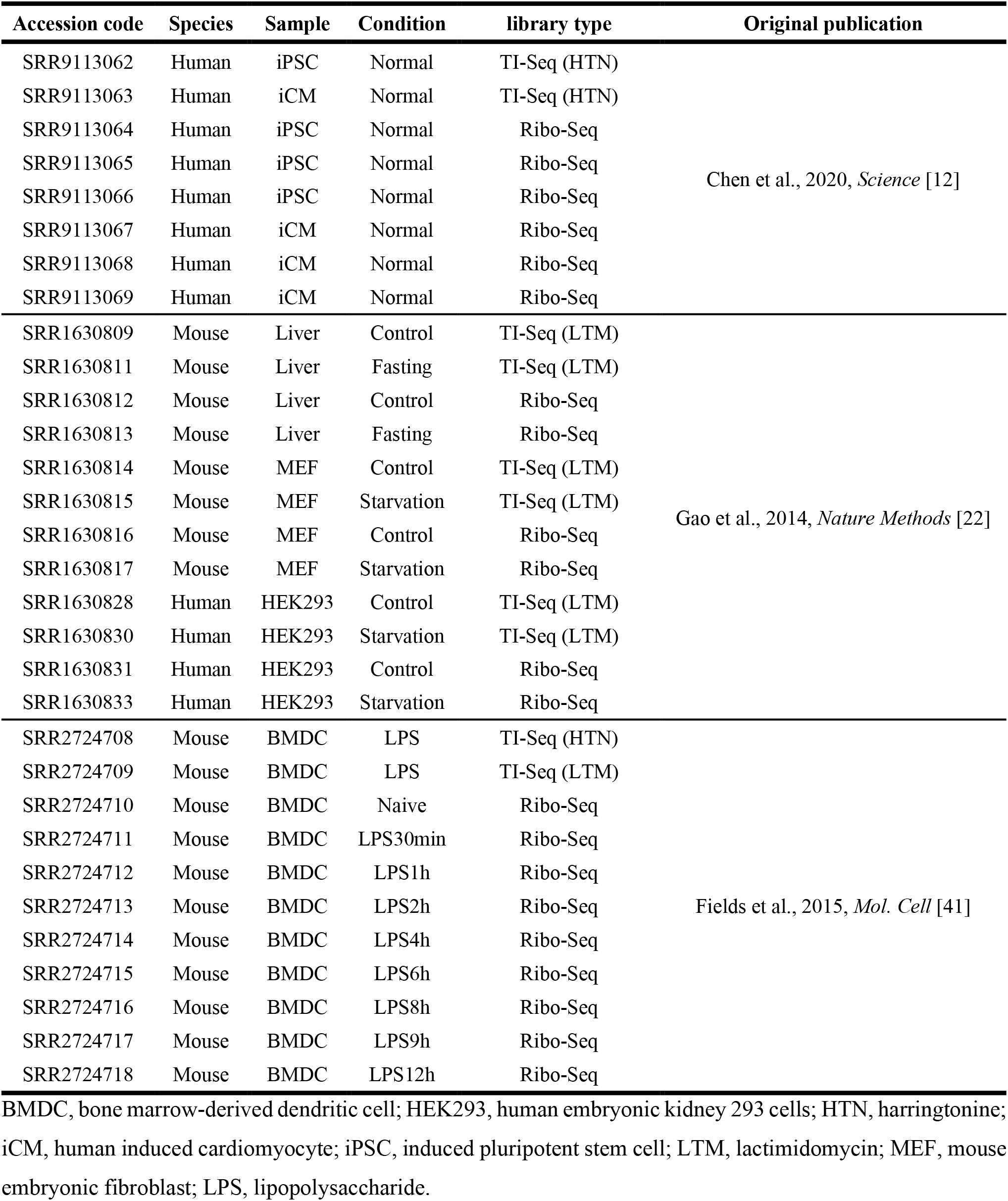
Description of the Ribo-Seq and matched TI-Seq datasets used in this study.

## Supplementary Figures

**Figure S1.**
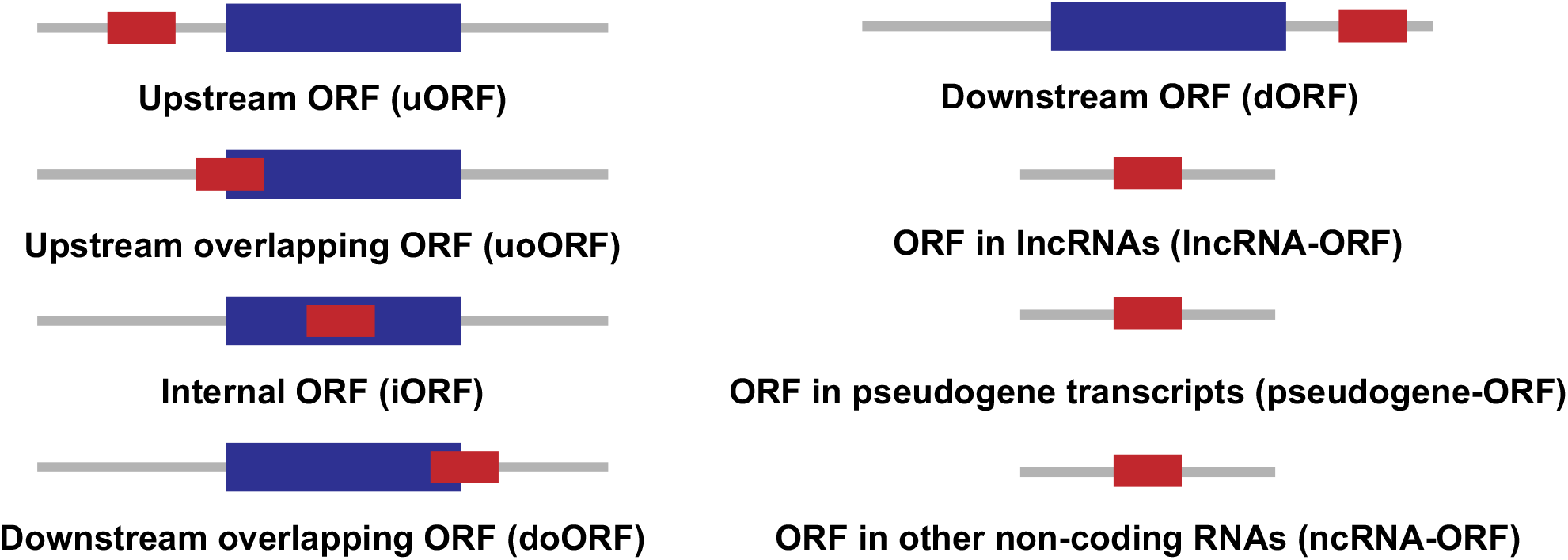
Illustration of different types of non-canonical ORFs.

**Figure S2.**
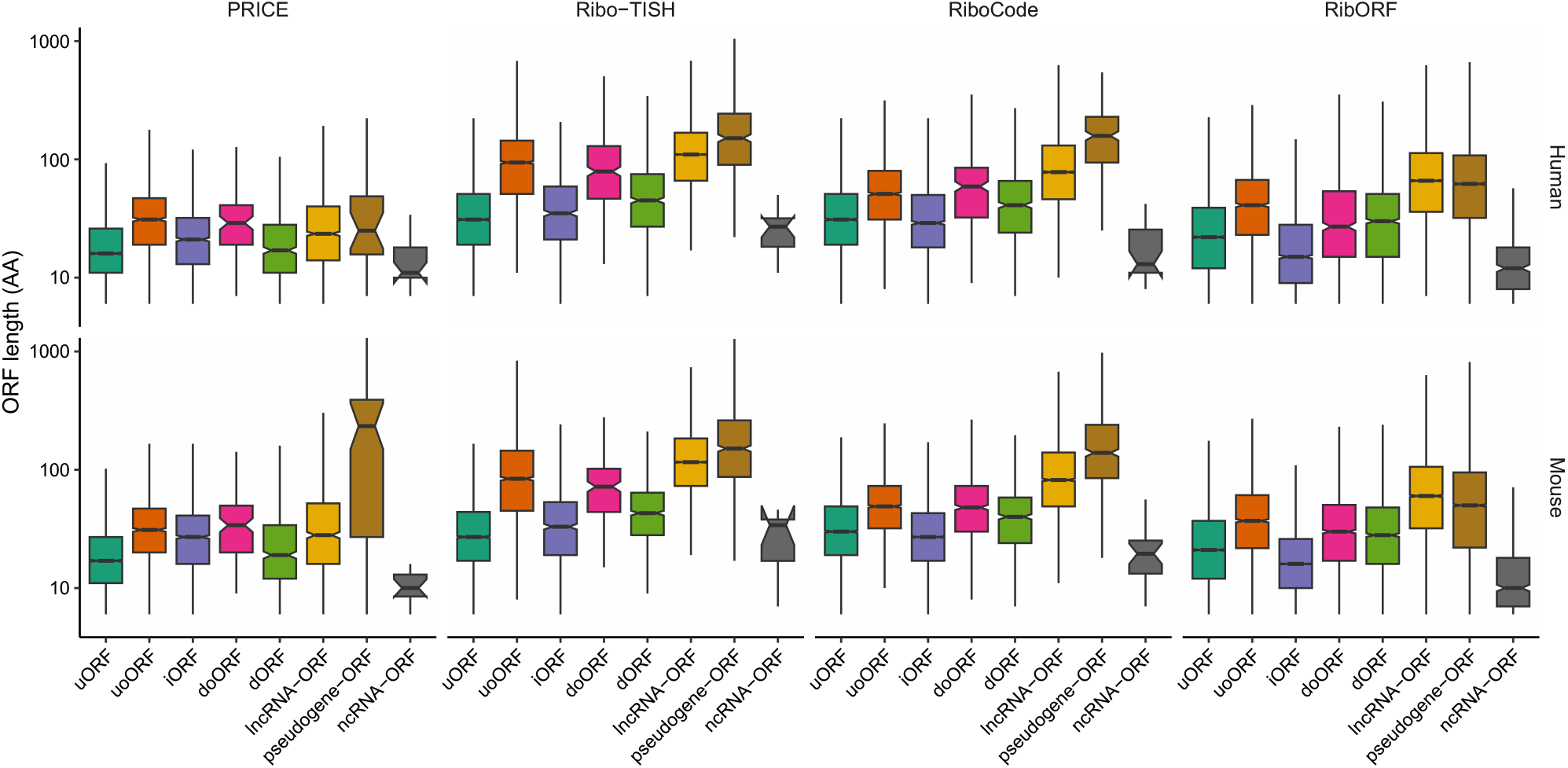
Length distributions of predicted non-canonical ORFs belonging to different types. For each species, ORFs predicted from different samples by the same method were pooled. Stop codons were included when calculating ORF lengths. AA, amino acid.

**Figure S3.**
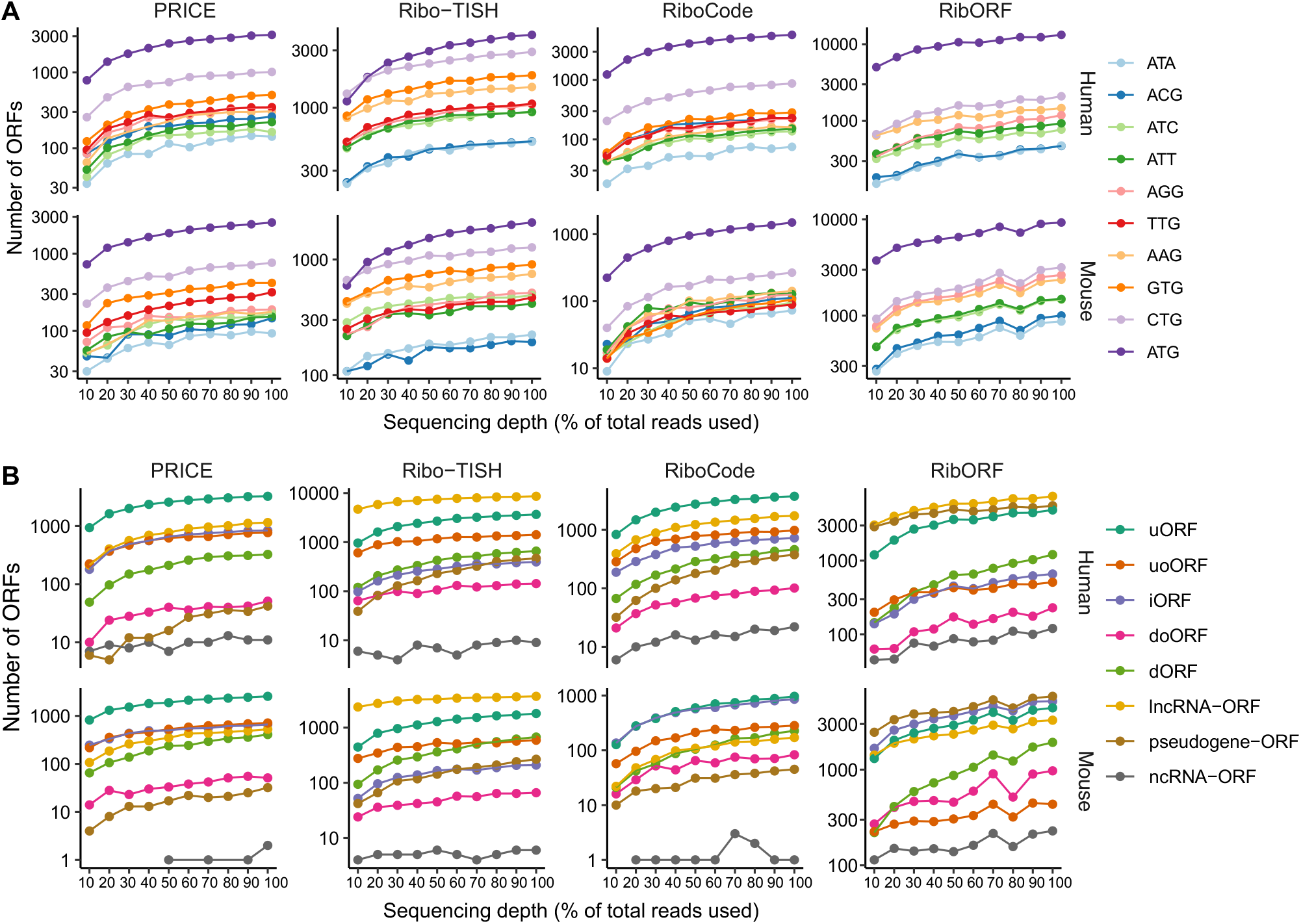
Number of detected non-canonical ORFs belonging to different categories (A) or with different start codons (B) when the different percent of RPF reads in the full data were used in the analysis.

**Figure S4.**
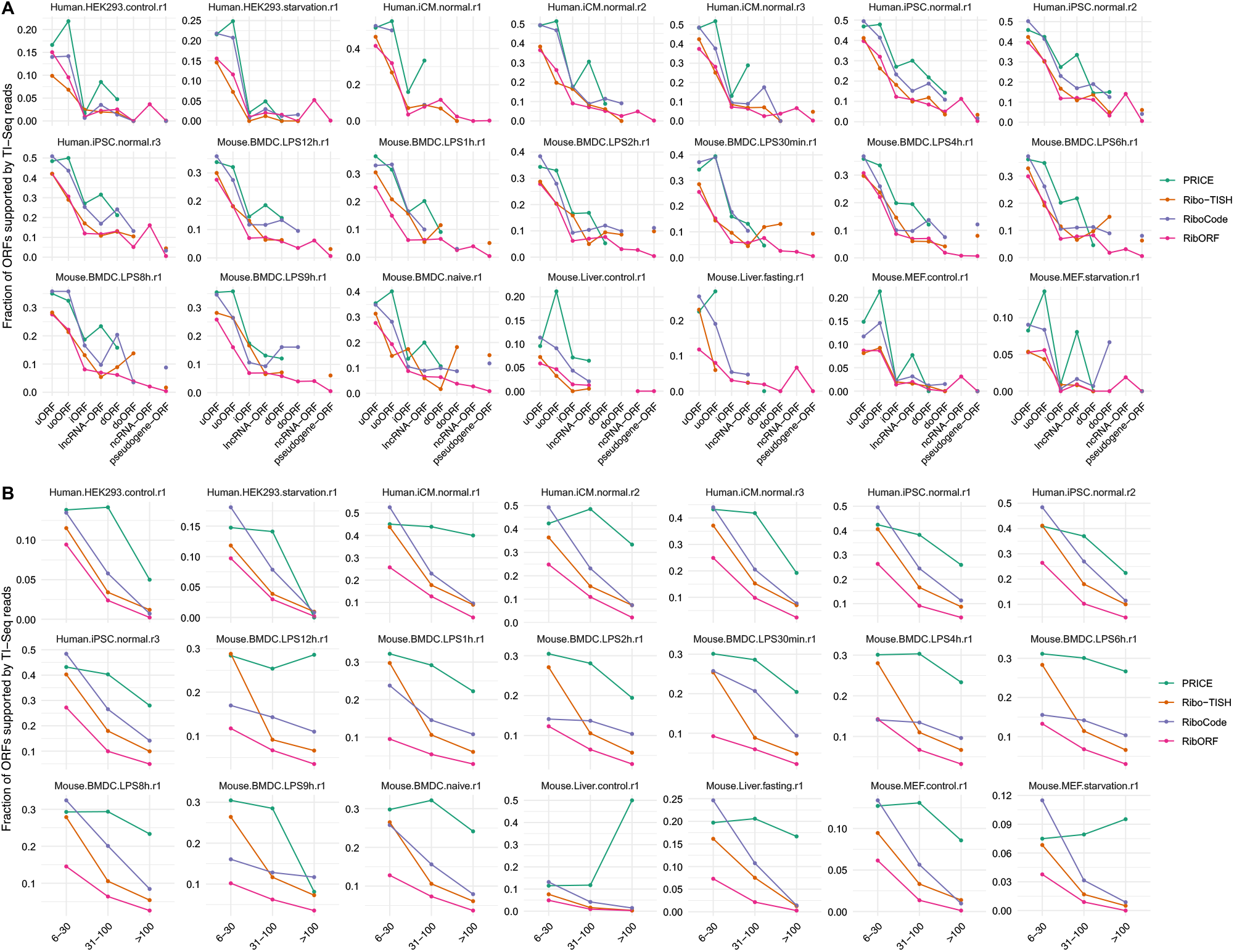
Proportions of ORFs belonging to different types (**A**) or of different lengths (**B**) that have peaks of RPF coverage around predicted start codons in matched TI-Seq libraries.

**Figure S5.**
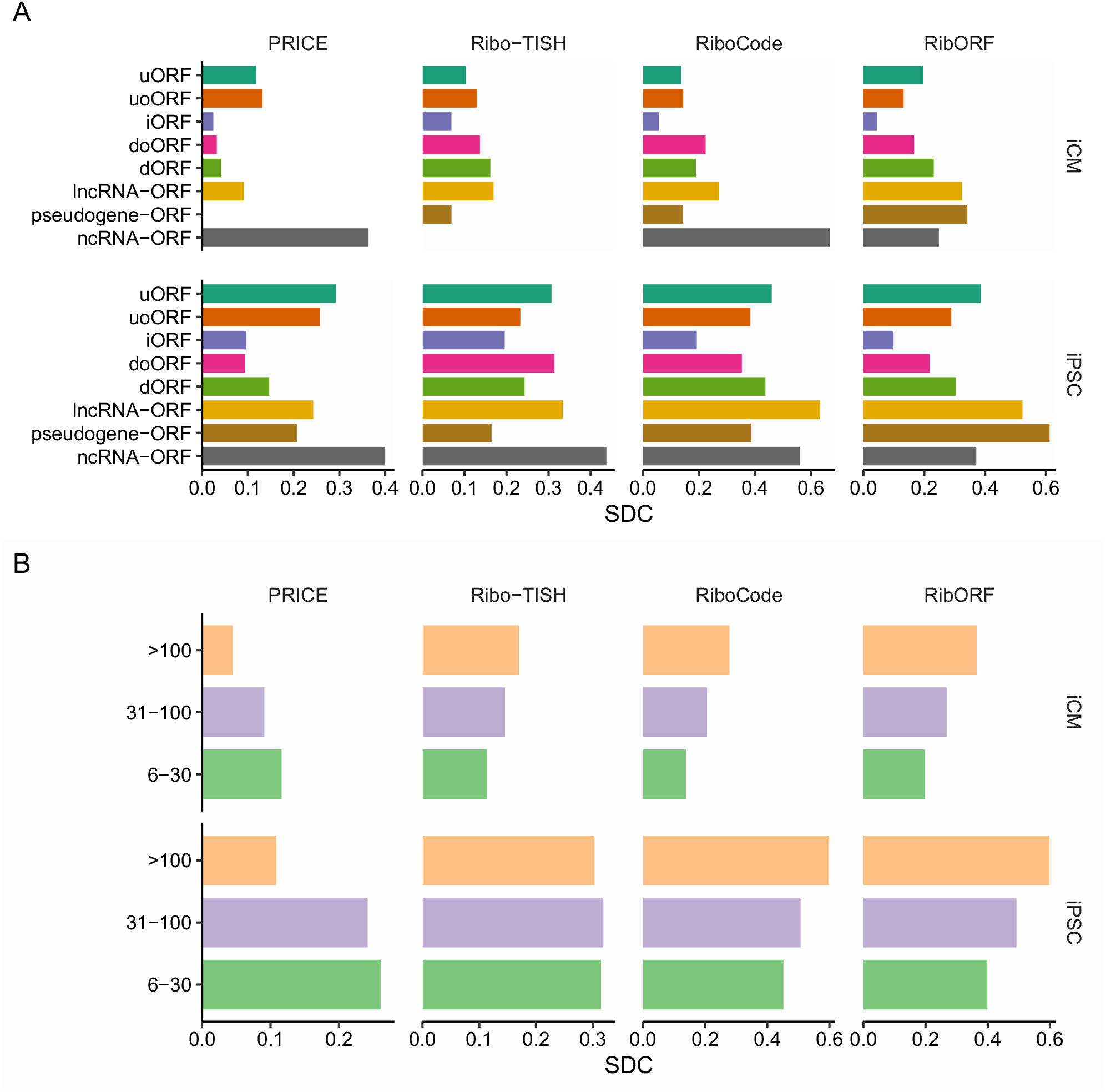
Similarity of predicted non-canonical ORFs across different biological replicates measured by Sørensen-Dice coefficient (SDC) and stratified by ORF type (**A**) or length (**B**).

**Figure S6.**
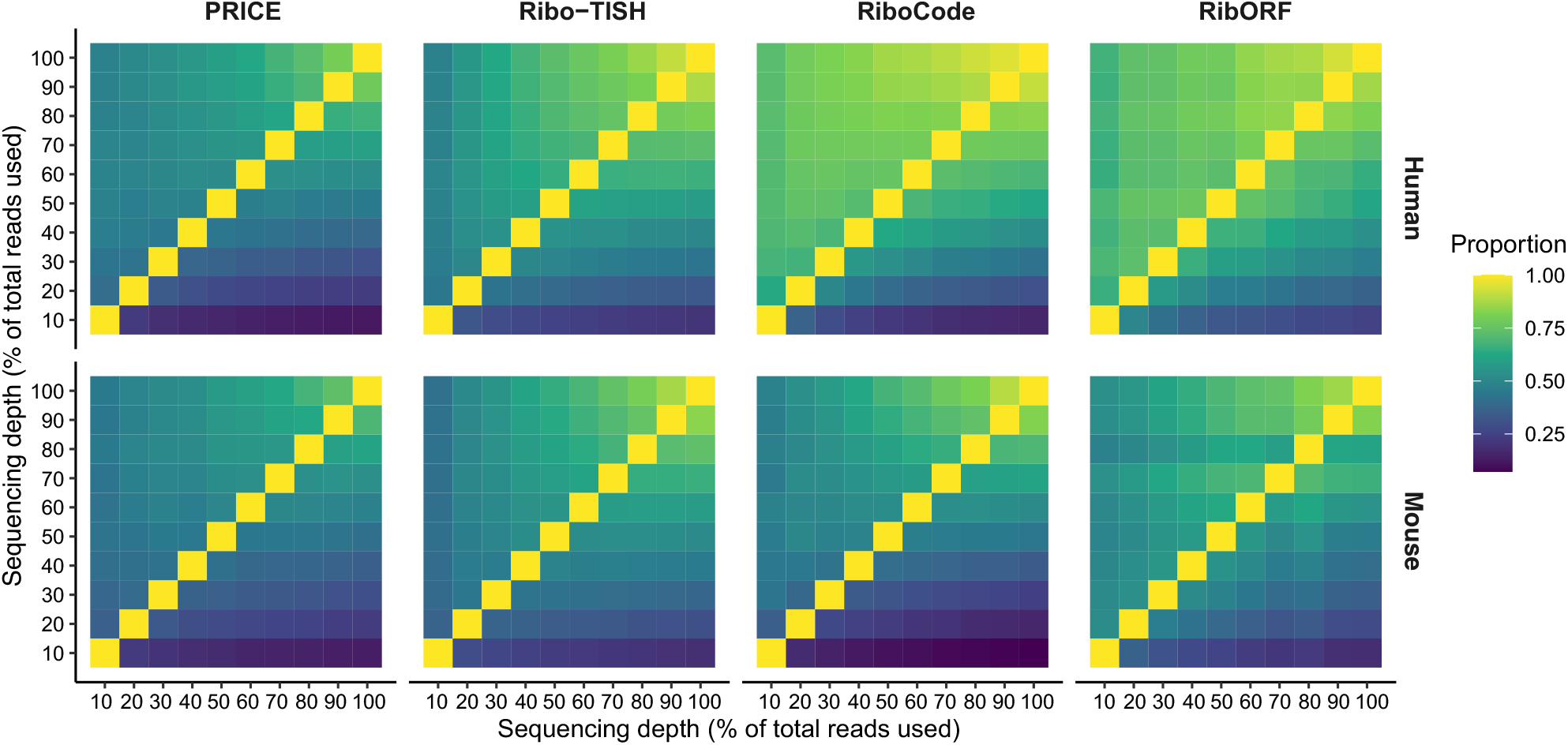
Proportion of non-canonical ORFs detected from a down-sampled dataset (*x*-axis) that are recovered in another down-sampled dataset (*y*-axis).

**Figure S7.**
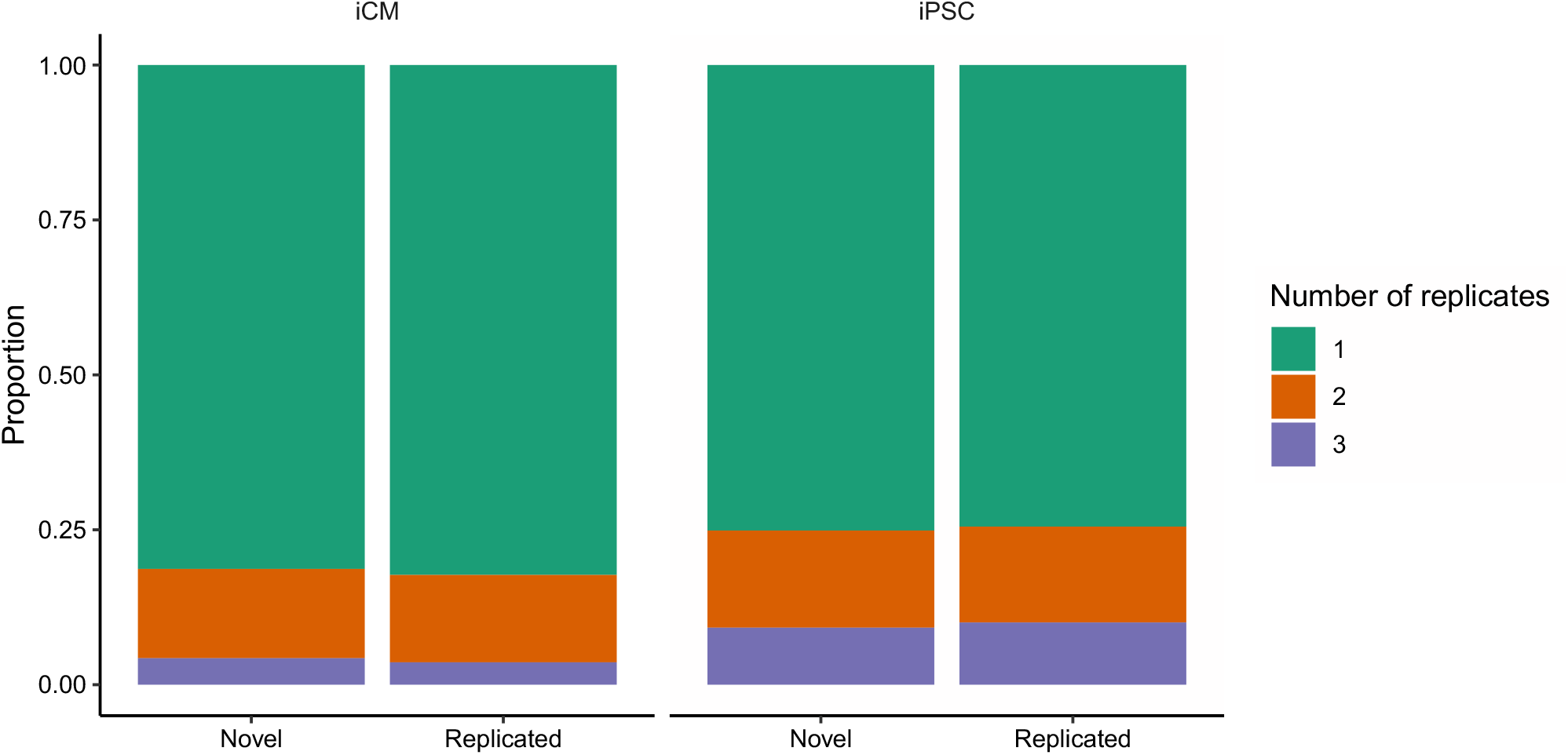
Proportion of ORFs that are consistently identified in different numbers of biological replicates for ORFs detected by single or multiple methods in each replicate of Ribo-Seq libraries of iCM or iPSC samples.

